# CRISPR/Cas9 mediated intersectional knockout of GSK3β in D2 receptor expressing mPFC neurons reveals contributions to emotional regulation

**DOI:** 10.1101/825166

**Authors:** Jivan Khlghatyan, Jean-Martin Beaulieu

## Abstract

**Background:** Glycogen synthase kinase 3β (GSK3β) regulates neurodevelopment, synaptic plasticity as well as mood, cognition, social interaction, and depressive-like behaviors. Inhibition of GSK3β is a shared consequence of treatment by lithium, SSRIs, ketamine and antipsychotics. GSK3β activity is regulated by dopamine D2 receptor signaling and can be inhibited by psychoactive drugs in a D2 receptor dependent manner. Functions of GSK3β in striatal D2 neurons has been studied extensively. However, GSK3β is ubiquitously expressed in the brain and D2 receptor expressing cells are distributed as a mosaic in multiple cortical regions. This complicates the interrogation of GSK3β functions in cortical D2 cells in a circuit defined manner using conventional animal models.

**Methods:** We have used a CRISPR/Cas9 mediated intersectional approach to achieve targeted deletion of GSK3β in D2 expressing neurons of the adult medial prefrontal cortex (mPFC).

**Results:** Isolation and analysis of ribosome associated RNA specifically from mPFC D2 neurons lacking GSK3β demonstrated large scale translatome alterations. Deletion of GSK3β in mPFC D2 neurons revealed its contribution to anxiety-related, cognitive, and social behaviors.

**Conclusions:** Our results underscore the viability of intersectional knockout approach to study functions of a ubiquitous gene in a network defined fashion while uncovering a contribution of GSK3β expressed in mPFC D2 neurons in the regulation of behavioral dimensions related to mood and emotions. This advances our understanding of GSK3β action at a brain circuit level and can potentially lead to the development of circuit selective therapeutics.

## Introduction

Gene products involved in disease processes and drug action are often expressed ubiquitously. Thus, their functional implication can be widely different in time (developmental stage) and space (brain region, neuronal circuit, particular cell type). Traditional approaches have allowed systemic as well as region or cell type-selective gene knockouts. However, their resolution remains limited and does not allow for the investigation of gene functions within cellular subpopulations belonging to the same brain region or neuronal circuit at a specific developmental stage.

Glycogen synthase kinase 3β (GSK3β) is a serine-threonine kinase that is ubiquitously expressed across the brain during the entire life span (1, 2). This kinase regulates neurodevelopment (2, 3), neuronal signaling and plasticity (4–7). GSK3β is also involved in regulating mood (8–11), cognition (12), social interaction and depressive-like behaviors (8, 12, 13). These functions of GSK3β have been identidied by using drugs that can systemically modulate its activity (14, 15) or by knockout or knockdown approaches, including germline heterozygous knockout (16), knockout only in forebrain CamKII expressing pyramidal neurons (8), Cre mediated knockout in adult prefrontal cortex (PFC) (9), CRISPR/Cas9 mediated knockout only in adult medial PFC (mPFC) neurons (11), shRNA mediated knockdown in adult nucleus accumbens shell neurons (13), and germline knockout in serotonergic neurons (17). However, the ubiquitous expression profile of GSK3β makes it difficult to precisely pinpoint the neuroanatomical correlates of these various functions.

Genome-wide association studies (GWAS) revealed that genetic variants of the dopamine D2 receptor (D2) are associated with depression (18, 19) and schizophrenia (20). D2 is a G-protein coupled receptor and a major target of antipsychotic drugs (21). This receptor has been shown to signal through Gαi/o to inhibit cAMP production (22, 23) and beta-arrestin-2 (βARR2) to inhibit AKT and activate GSK3β (24). Treatment with selective serotonin reuptake inhibitors (SSRIs) and ketamine also inhibit the activity of GSK3β (15, 25, 26). Furthermore, GSK3β activity can be inhibited by mood stabilizers in a D2 and βARR2 dependent manner (9, 14, 27). A study involving germline knockout of GSK3β in D2 expressing neurons indicated that GSK3β mediates the locomotor response to lithium and antipsychotics (28), most likely by interfering with D2 signaling in striatal medium spiny neurons. However, the potential contribution of GSK3β in the regulation of other behavioral dimensions by D2 expressing neurons of other brain regions remains unknown.

We have recently identified multiple cortical regions with a mosaic distribution of D2 expressing neurons, ranging from 5-50% of total neurons within a given region (29). This raises questions about the role of GSK3β in D2 expressing neurons of these various cortical regions. However, it has been technically challenging to achieve knockout of GSK3β in brain region-specific and cell type-specific manner at a given developmental stage. In other words, it has been impossible to eliminate GSK3β expression in a given brain region while targeting only D2 neurons. Considering the involvement of mPFC in mental disorders (18, 30) and the presence of D2 expressing neurons in this region (29) knocking out GSK3β in adult mPFC D2 neurons represents an ideal model to explore approaches involving high-resolution gene targeting in a specific cellular subset belonging to brain region composed of a heterogeneous neuronal population.

We used an intersectional approach involving the combination of single guide RNA (sgRNA) viral delivery with Cre mediated conditional Cas9 expression to target GSK3β only in mPFC D2 neurons of adult mice. Combination of this approach with a conditional RiboTag reporter system allowed to extract ribosome associated RNA specifically from mPFC D2 neurons and demonstrated how GSK3β KO affects neuronal translatome. Moreover, knockout of GSK3β in mPFC D2 neurons uncovered its involvement in the regulation of anxiety-related, cognitive, and social behaviors.

## Materials and Methods

Detailed Materials and Methods are described in Supplementary material

### Animals

D2Cre heterozygous bacterial artificial chromosome (BAC) transgenic mice (31) (GENSAT RRID: MMRRC_017263-UCD) were crossed to heterozygous Rosa26-LSL-Cas9 knockin mice (Stock No: 024857, The Jackson Laboratory) to generate D2Cre/LSL-Flag-Cas9 line. 2.5-3 months old male mice from this line were used for virus injections, immunohistochemistry, and behaviors.

Homozygous knockin RiboTag mice (32) were crossed to D2Cre/LSL-Flag-Cas9 mouse line to generate D2Cre/LSL-Flag-Cas9/RiboTag mice. 2.5-3 months old male mice from this line were used for viral injections followed by ribosome-associated mRNA isolation and RNAseq.

Mice were maintained on a 12-hours light/dark cycle with ad libitum access to food and water. All experiments conducted in this study are approved by Université Laval and University of Toronto Institutional Animal Care Committee in line with guidelines from the Canadian Council on Animal Care.

### Mouse stereotaxic surgery and AAV viruses

Mice were anesthetized with a preparation of ketamine 10mg/ml and xylazine 1mg/ml (0.1ml/10g, i.p.). The animal was placed in a stereotaxic frame, and the skull surface was exposed. Two holes were drilled at injection sites and virus (AAV- GFP or AAV-Gsk3sgRNA/GFP) was injected using an injector with a microsyringe pump controller (WPI) at the speed of 4nl per second. AAV- GFP and AAV-Gsk3sgRNA/GFP viral particles were described previously (11). Following injection coordinates for mPFC were used: AP +2.4, ML±0.5, DV −1.7. All measures were taken before, during, and after surgery to minimize animal pain and discomfort.

### Immunohistochemistry and quantification

Mice were euthanized 3 weeks after viral delivery by a lethal dose of ketamine/xylazine and perfused with phosphate buffer saline (PBS) followed by 4% paraformaldehyde (PFA). Brains were incubated in 4% PFA 24h at 4°C. Fixed tissue was sectioned using vibratome (Leica, VT1000S). Flag-Cas9 staining required antigen retrieval. Thus, 40 μm sections had to be boiled for 2 min in sodium citrate buffer (10 mM tri-sodium citrate dehydrate, 0.05% Tween-20, pH 6.0) and cooled down at room temperature (RT) for 20 min. This resulted in the disappearance of endogenous GFP signal in D2Cre/LSL-Flag-Cas9 mice (see Supplementary Figure 1c). However, virally expressed GFP signal was still present (Figure 2). Sections were blocked and permeabilized with a permeabilization solution containing 10% normal goat serum (NGS) and 0.5% Triton X-100 (Sigma) in PBS for 2 h. Sections were incubated with primary antibodies diluted in permeabilization solution overnight at 4 °C. After three washes in PBS, samples were incubated with secondary antibodies for 2h at room temperature. After washing with PBS three times, sections were mounted using DAKO mounting medium (DAKO, Mississauga, Canada) and visualized with a confocal microscope (Zeiss LSM 880, Zen 2011 Software, Oberkochen, Germany).

For quantification every 3^rd^ serial coronal section was taken as shown in Supplementary Figure 1b. In every region of interest (for example prelimbic layer 5 or Cingulate Layer 5) 3 randomly chosen Z stack pictures were taken with a 60x magnification objective. Quantification was performed manually, using ImageJ (National Institute of Health (NIH), Bethesda, MD).

Following primary antibodies were used: mouse anti-Flag (1:500, Sigma-Aldrich F1804), rabbit anti-Gsk3β (1:500, Cell Signal Technology 9315, Danvers, MA).

Following secondary antibodies were used: Goat anti-mouse Alexa 647, Goat anti-rabbit Alexa 568 (1:1000, Invitrogen).

Cell line culture, transfection and western blot were described before (11). Behavioral tests, Z scoring was performed as described before (8, 33−36). Tissue dissection, Immunoprecipitation of polyribosomes and RNA isolation were described before (29)

### Drugs

30mg/kg methylphenidate was dissolved in saline and given to mice intraperitoneally (i.p.).

### Tissue dissection, Immunoprecipitation of polyribosomes and RNA isolation

Mice were killed by rapid cervical dislocation. Heads of animals were immediately cooled by immersion in liquid nitrogen for 6 seconds. First, 500um thick serial coronal sections were prepared using ice-cold adult mouse brain slicer and matrix (Zivic instruments), second, mPFC was dissected on ice cold surface using a microsurgical knife (KF Technology). Immunoprecipitation of polyribosomes was performed as described before ^4^. Tissue samples were lysed in homogenization buffer (50mMTris, pH 7.5, 100 mM KCl, 12 mM MgCl2, 1% Nonidet P-40, 1 mM DTT, 100U/mL RNase Out, 100 μg/mL cycloheximide, Sigma protease inhibitor mixture) followed by centrifugation for 10 min at 10000g. Anti-hemagglutinin (HA) antibody (1:150; MMS-101R; BioLegend) was added into collected supernatant and tubes were kept under constant rotation for 4 hours at 4°C. Protein G magnetic beads (Life Technologies) were washed 3 times with homogenization buffer then added into the mixture and kept for constant rotation overnight at 4°C. The following day magnetic beads were washed three times with high salt buffer (50mMTris, pH 7.5, 300 mM KCl, 12 mM MgCl2, 1% Nonidet P-40, 1 mM DTT, 100U/mL RNase Out, 100 μg/mL cycloheximide, Sigma protease inhibitor mixture). RNA was extracted by adding TRI reagent (Zymo research) to magnetic beads pellet followed by Direct-zol RNA kit according to the manufacturer’s instructions (Zymo research). The RNA concentration was quantified using ND-1000 Spectrophotometer (NanoDrop Technologies).

For RNAseq analysis RNA was isolated from 9 mice per group. Each group consisted of 3 biological replicates and each replicate represented RNA pooled from 3 mice.

### RNAseq analysis

#### Quality control

The quality control metrics for the RNA-seq data were obtained using the tool RNA-SeQC (v1.1.7). For more information, please visit their website found here: http://www.broadinstitute.org/cancer/cga/rna-seqc. This program takes aligned files as input and delivers a series of plots and statistics for each sample. Based on the output for each sample, the RNA sequencing quality was deemed acceptable for further analysis.

#### Processing Pipeline

All raw FASTQ files were aligned to the appropriate mouse genome (GRCm38) using the HISAT2 aligner. HISAT2 is a fast and sensitive alignment program that uses a large set of small graph FM (GFM) indexes that collectively cover the whole reference genome. These local indexes, in conjunction with a series of alignment strategies, ensure a rapid and accurate alignment of sequencing reads. Accessory programs for the alignment stage include SAMTOOLS (v1.3.1) and BEDTOOLS (v2.26.0). Alignment files were sorted by their genomic location and indexed using SAMTOOLS. These sorted binary SAM (BAM) files were then used as input for StringTie (v1.3.4), which assembles RNA-seq alignments into potential transcripts. It uses a novel network flow algorithm as well as an optional de novo assembly step to assemble and quantitate full-length transcripts representing multiple splice variants for each gene locus. Finally, in order to identify differentially expressed genes between samples, the Ballgown R-package was implemented (v3.4.3). Transcript-level FPKMs were estimated using Tablemaker. Expression was estimated for each transcript, exon, and intron (junction) in the assembly. All of the statistical analysis (organization, visualization, etc.) was conducted with the tools available within the Ballgown package.

#### Differential Expression Analysis

The statistical test applied to this data was a parametric F-test comparing nested linear models; details are available in the Ballgown manuscript. Briefly, two models are fit to each feature, using the expression as the outcome: one including the covariate of interest (e.g., case/control status) and one not including that covariate. An F statistic and p-value are calculated using the fits of the two models. A significant p-value means that the model including the covariate of interest fits significantly better than the model without that covariate, indicating differential expression. All the differentially expressed transcripts (DETs) with p < 0.05 were selected for further analysis. Differential expression testing was carried out for Ctrl vs Gsk3sKOinD2 comparison.

Heat map of gene expression Z score (Fig. 3) was generated using Morpheus tool (https://software.broadinstitute.org/morpheus/). The Z score for gene expression was calculates using the formula: Z = (X−μ)/σ, which indicates how many standard deviations (σ) the level of expression of a given gene (X) is above or below the average expression from all the samples (μ).

#### Gene set enrichment analysis

First DETs were filtered. All selected transcripts had a mean expression >0.5 FKPM. Fold change (FC) threshold was set to 0.7<FC>1.3. Enrichment analyses and visualization were performed following the pipeline described in Reimand et al ^5^. Enrichment analysis was performed using gProfiler (https://biit.cs.ut.ee/gprofiler/gost). Term size for pathways was selected to be min 5 and max 150. Only pathways passing significance threshold of 0.05 were selected. Gene enrichment in Gene Ontology Biological pathways (GO:BP) were selected. Then .GEM and .GMT files were downloaded from gProfiler and were used in EnrichmentMap app of Cytoscape (Ver 3.7.1) for visualization. Following parameters of EnrichmentMap were used: FDR q value cutoff <0.05, Jaccard combined > 0.375, overlap > 0.5 and Prefuse Force Directed layout was chosen. Then ClusterMaker2 App was used to cluster enriched pathways based on similarity and AutoAnnotate and WordCloud Apps were used to name clusters of enriched pathways using default parameters.

The final pictures of the clusters of enriched pathways are shown in Fig. 3 and Supplementary Figure 2. The list of all the enriched pathways is shown in Supplementary Table 2.

### Data analysis and statistics

Data are presented as means ± SEM. Two-tailed t test is used in GraphPadPrism 5 software for comparison between two groups (La Jolla, CA) (*p < 0.05, **p < 0.01, ***p < 0.001).

## Results

### Efficient and specific knockout of GSK3β by CRISPR/Cas9

To investigate the efficacy and specificity of CRISPR/Cas9 mediated knockout of GSK3β, we tested the ability of previously characterized *Gsk3b* sgRNA (11) to cut and induce mutations of DNA at on- and off-target sites. Targeting efficacy and specificity of CRISPR/Cas9 system depends on the sequence of the guide RNA (37, 38). It has been shown that genomic regions that contain few mismatches to the guide RNA sequence can represent potential off-target sites (38). We have used two previously characterized algorithms (38, 39) to predict putative off-target sites. Both algorithms showed that mouse genome does not contain sites with one or two mismatches to *Gsk3b* sgRNA. We have identified only three putative off-target sites containing three mismatches (Figure 1B-D). All three putative off-target sites were positioned in intergenic regions with no coding activity (Figure 1B-D). Sites with four and more mismatches were disregarded, since predicted activity of sgRNA at those sites was virtually none (37, 38). We transfected mouse Neuro2A cells with an “all in one” CRISPR plasmid containing previously characterized *Gsk3b* sgRNA(11) or scrambled sgRNA as a control. Genomic DNA was isolated and tracking of indels by decomposition (TIDE) analysis (40) was applied to find activity at on-target (Gsk3b Exon 3) and off-target genomic loci (Figure 1A-D). At on-target site, in Ctrl condition no insertions or deletions were detected and 100% of DNA sequences were not mutated (position “0”) (Figure 1A). At on-target site, in *Gsk3b* sgRNA condition multiple deletions and insertions were detected with one nucleotide insertion accounting for 47% of mutated DNA sequences (position “+1”) (Figure 1A). Overall, only 3.5% of DNA sequences were not mutated (position “0”) (Figure 1A). At all three of off-target sites virtually no mutations were detected in both Ctrl and *Gsk3b* sgRNA conditions (Figure 1B-D).

**Figure 1.**
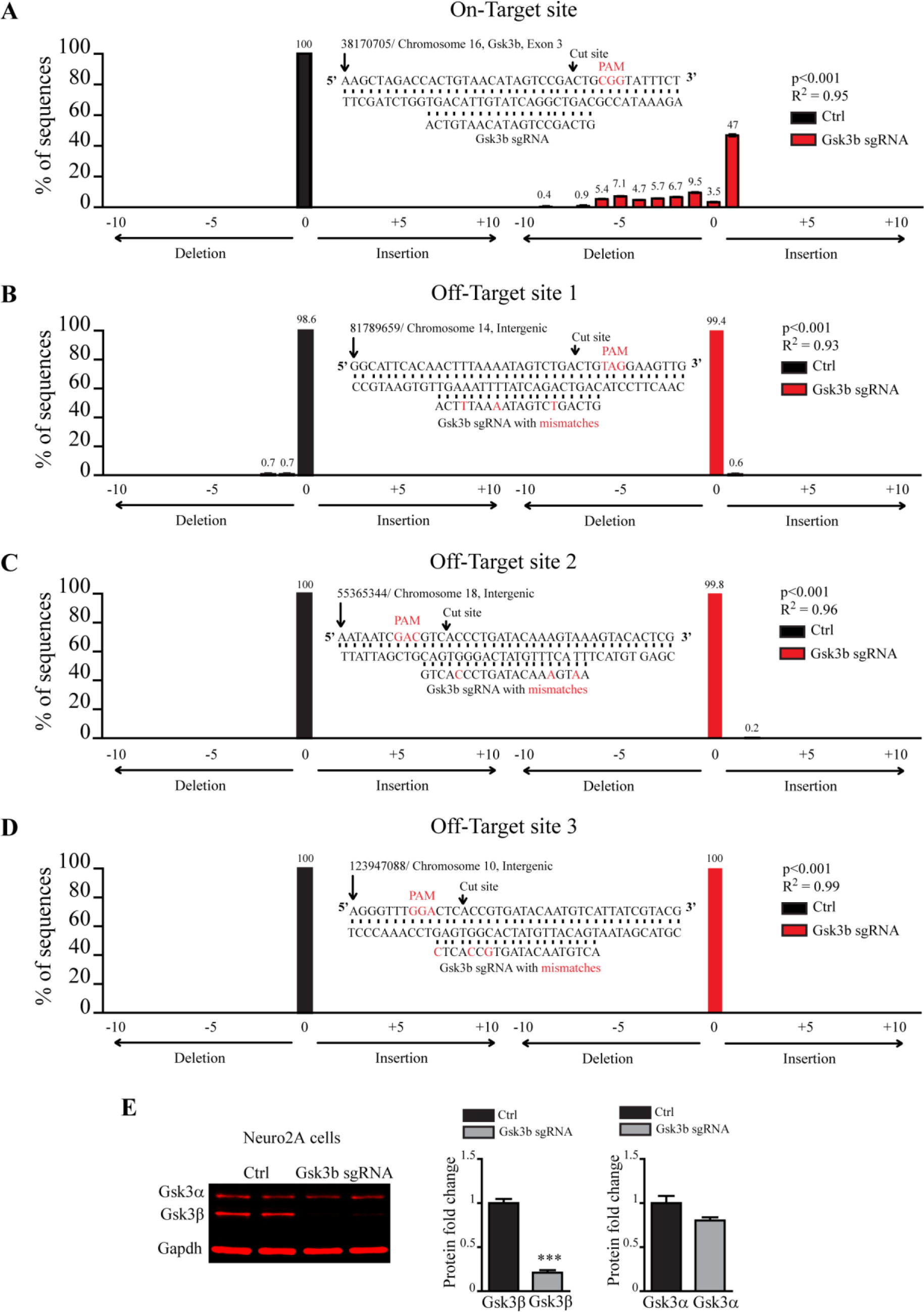
Specificity of CRISPR/Cas9 mediated Gsk3β knockout. (A-D) Quantification of mutations using TIDE analysis at (A) on-target and (B-D) three putative off-target sites. Position “0” represents % of non mutated sequence, “−” positions are deletions and “+” positions are insertions. Middle panels show nucleotide sequence and chromosomal position at a putative target site, Gsk3b sgRNA as well as mismatch nucleotides at off-target sites. (E) Quantification of the expression of Gsk3α and Gsk3β after CRISPR/Cas9 mediated knockout of Gsk3β in Neuro2A cells (Gsk3α: Ctrl 1± 0.08 n=5, Gsk3b sgRNA 0.8 ± 0.035 n=4, Gsk3β: Ctrl 1 ± 0.047 n=5, Gsk3b sgRNA 0.21 ± 0.028 n=5, *p < 0.05, Student T-test). Left panel: an example of western blot membrane stained for Gsk3α, Gsk3β, and Gapdh.

Mutation in the exon of *Gsk3b* can disrupt its protein expression. Indeed western blot analysis showed a dramatic reduction of GSK3β expression in *Gsk3b* sgRNA transfected cells as compared to controls, while GSK3α expression did not differ between conditions (Figure 1E, data and statistics are indicated in figure legends). Overall, this shows that CRISPR/Cas9 mediated knockout of GSK3β is robust and specific.

### Efficient knockout of GSK3β in mPFC D2 neurons of adult mice

To achieve somatic knockout of GSK3β exclusively in D2 expressing mPFC neurons (Gsk3sKO in mPFC D2) first, expression of Cas9 was activated only in D2 cells by crossing D2Cre mice (31) with LSL-Flag-Cas9 mice (41). Second, *Gsk3b* sgRNA containing virus (AAV Gsk3 sgRNA/GFP) (11) or AAV GFP (as a control) were injected into the mPFC of these mice (Figure S1A). The selectivity of this D2Cre mouse line was demonstrated previously with Cre expression in Prelimbic (PL) layer 2,3 and layer 5, Infralimbic (IL) layer 2,3 and layer 5 and Cingulate (Cg) layer 5 of adult mouse mPFC (29).

Three weeks after injection of AAV Gsk3 sgRNA/GFP) (11) or AAV GFP viruses the infection area included PL, IL and Cg cortices as shown by the expression of GFP in serial coronal sections of mouse brain (Figure S1B). Thus we investigated the ability of intersectional CRISPR/Cas9 to knockout GSK3β in D2 neurons of PL, IL and Cg cortices. Flag-Cas9 staining indicated the presence of Cas9 and virally expressed GFP signal indicated the presence of *Gsk3b* sgRNA (see Materials and Methods and Figure S1C). Vast majority of neurons containing Flag-Cas9 and *Gsk3* sgRNA (infected with AAV Gsk3 sgRNA/GFP) were devoid of GSK3β signal, while GSK3β staining was always present in neurons that only express GFP (infected with AAV Gsk3 sgRNA/GFP) (Figure 2A-F and Figure S1D,E). To determine the extent of the knockout, we also quantified % of GSK3β expressing flag-Cas9 cells out of all flag-Cas9 cells per brain region. Again, vast majority of flag-Cas9 cells were devoid of GSK3β staining, indicating widespread knockout (Figure 2A-F and Figure S1D,E). Neurons infected with AAV GFP express GSK3β regardless of the presence of Flag-Cas9 (Figure 2A-F and Figure S1D,E). Moreover, flag-Cas9 neurons in Striatum of both AAV Gsk3 sgRNA/GFP or AAV GFP infected mice also express GSK3β (Figure 2F, Figure S1F), since these neurons were not infected by viruses.

**Figure 2.**
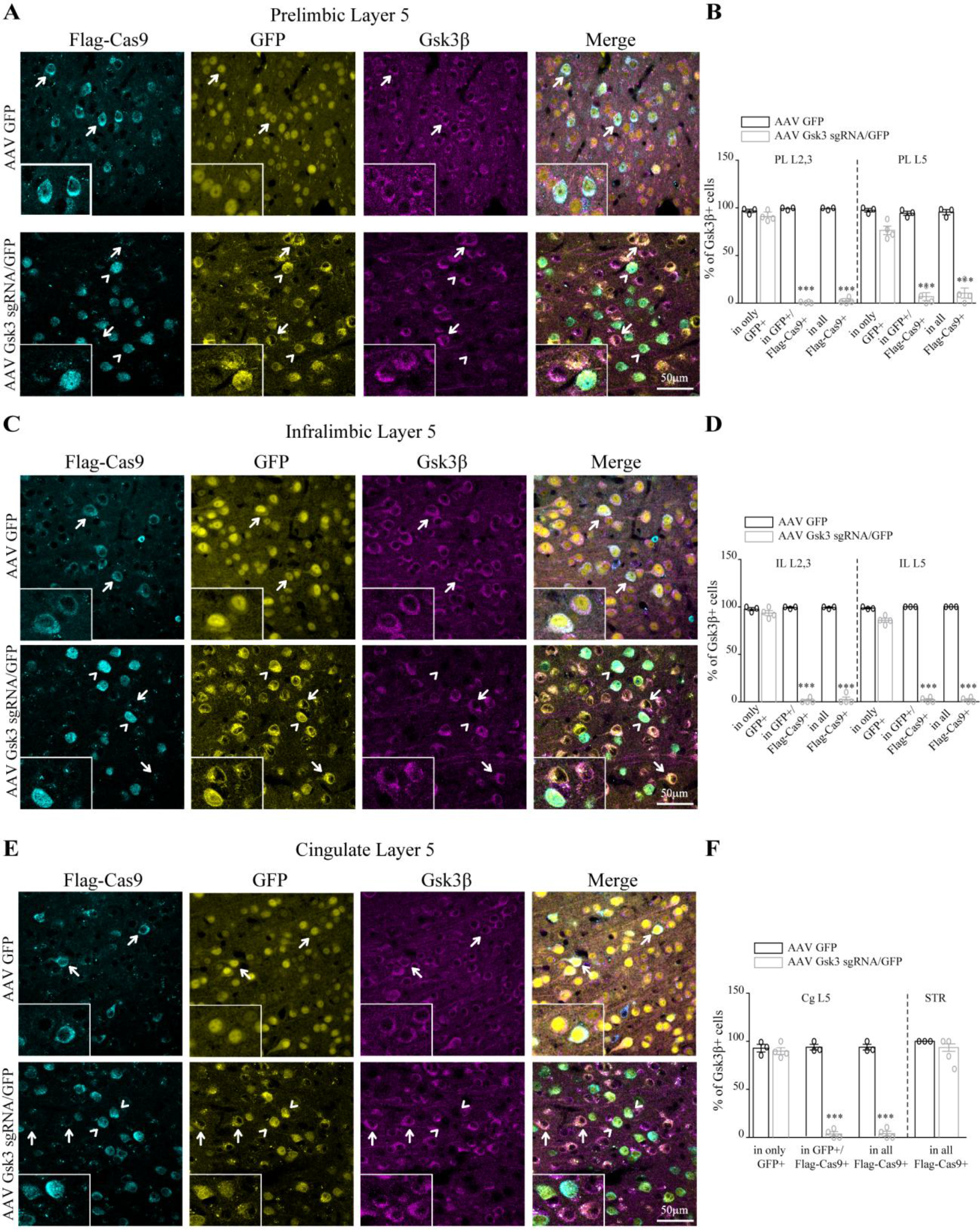
Knockout of GSK3β in D2 neurons of mPFC. (A) Immunofluorescent staining for Flag-Cas9 and Gsk3β in Prelimbic layer 5 of virus injected D2Cre/LSL-Flag-Cas9 mice. Insets show the zoomed picture. (B) Quantification of the % of Gsk3β expressing cells (PL L2,3 AAV GFP condition: in only GFP 96.8±1.7, in GFP+/flagCas9+ 99.2±0.75, in all flagCas9+ 99.2±0.75 n=3. AAV Gsk3 sgRNA/GFP condition: in only GFP 90.9±2.5, in GFP+/flagCas9+ 0.6±0.36, in all flagCas9+ 2.2±1.4 n=4, PL L5 AAV GFP condition: in only GFP 97.5±1.7, in GFP+/flagCas9+ 94.3±2.2, in all flagCas9+ 95.5±2.6 n=3. AAV Gsk3 sgRNA/GFP condition: in only GFP 76±5.4, in GFP+/flagCas9+ 9±4.1, in all flagCas9+ 14±4.8 n=4 ***p<0.0001, one way ANOVA). (C) Immunofluorescent staining for Flag-Cas9 and Gsk3β in Infralimbic layer 5 of virus injected D2Cre/LSL-Flag-Cas9 mice. Insets show the zoomed picture. (D) Quantification of the % of Gsk3β expressing cells (IL L2,3 AAV GFP condition: in only GFP 97.7±1.6, in GFP+/flagCas9+ 99.3±0.6, in all flagCas9+ 99.3±0.6 n=3. AAV Gsk3 sgRNA/GFP condition: in only GFP 92.2±1.7, in GFP+/flagCas9+ 1.6±1.4, in all flagCas9+ 3.4±2.9 n=4, IL L5 AAV GFP condition: in only GFP 98.6±0.6, in GFP+/flagCas9+ 100±0, in all flagCas9+ 100±0 n=3. AAV Gsk3 sgRNA/GFP condition: in only GFP 87.5±1.5, in GFP+/flagCas9+ 1.6±1.4, in all flagCas9+ 1.6±1.4 n=4 ***p<0.0001, one way ANOVA). (E) Immunofluorescent staining for Flag-Cas9 and Gsk3β in Cingulate layer 5 of virus injected D2Cre/LSL-Flag-Cas9 mice. Insets show the zoomed picture. (F) Quantification of the % of Gsk3β expressing cells (Cg L5 AAV GFP condition: in only GFP 92.9±4, in GFP+/flagCas9+ 93.9±3, in all flagCas9+ 93.9±3 n=3. AAV Gsk3 sgRNA/GFP condition: in only GFP 90.6±4.2, in GFP+/flagCas9+ 3.1±2.6, in all flagCas9+ 3.5±3 n=4, STR AAV GFP condition: in all flagCas9+ 100±0 n=3. AAV Gsk3 sgRNA/GFP condition: in all flagCas9+ 1.6±1.4 n=4 ***p<0.0001, one way ANOVA). Error bars show standard error of the mean (SEM). Note that in AAV GFP injected control condition all cells express Gsk3β (indicated by arrows). In AAV Gsk3 sgRNA/GFP injected condition only cells that express Flag-Cas9 (corresponding to D2 cells) and sgRNA/GFP do not have Gsk3β signal (indicated by arrowheads), while cells having sgRNA/GFP, but not Flag-Cas9 staining (not D2 cells) still express Gsk3β (indicated with arrows).

Overall, this demonstrates that we can achieve highly efficient somatic knockout of GSK3β exclusively in D2 neurons of the adult mPFC without affecting neighboring cells and D2 neurons in other brain regions.

### Gsk3sKO in mPFC D2 neurons alters translatome

We used a Cre activated RiboTag reporter system and RNAseq to investigate the impact of GSK3β knockout on the translatome of mPFC D2 cells. To isolate translatome specifically from mPFC D2 neurons, we crossed D2Cre/LSL-Flag-Cas9 mice with RiboTag mice (32). D2Cre/LSL-Flag-Cas9/RiboTag mice expressed Cas9 and HA-tagged Rpl22 ribosomal subunit only in D2 neurons. The D2Cre/RiboTag system has been shown to be robust and specific for the isolation of ribosome-bound RNA from D2 cells (29). AAV Gsk3sgRNA/GFP (Gsk3sKO in mPFC D2) or AAV GFP (Ctrl) was injected into mPFC of D2Cre/LSL-Flag-Cas9/RiboTag mice. mPFC were dissected three weeks after viral infection. RiboTag-immunoprecipitation and RNA extraction was performed, followed by RNAseq (Figure 3A). We identified 1145 unique differentially expressed transcripts (DETs) in Gsk3sKO in mPFC D2 compared to Ctrl (Figure 3B, Table S1). To identify biological dimensions based on DETs, we performed biological pathway enrichment (GO:BP) using gProfiler and clustering using EnrichmentMap and ClusterMaker in Cytoscape (Figure 3C, Figure S2, Table S2) (42). We then manually separated enriched clusters into “Neuronal” and “General” functions. The biggest and most enriched “Neuronal function” clusters were related to postsynaptic structure and function as well as neurotransmission (Figure 3C). The biggest and most enriched “General function” clusters were Purine metabolism, Ubiquitine dependent catabolism and Histone acetylation.

**Figure 3.**
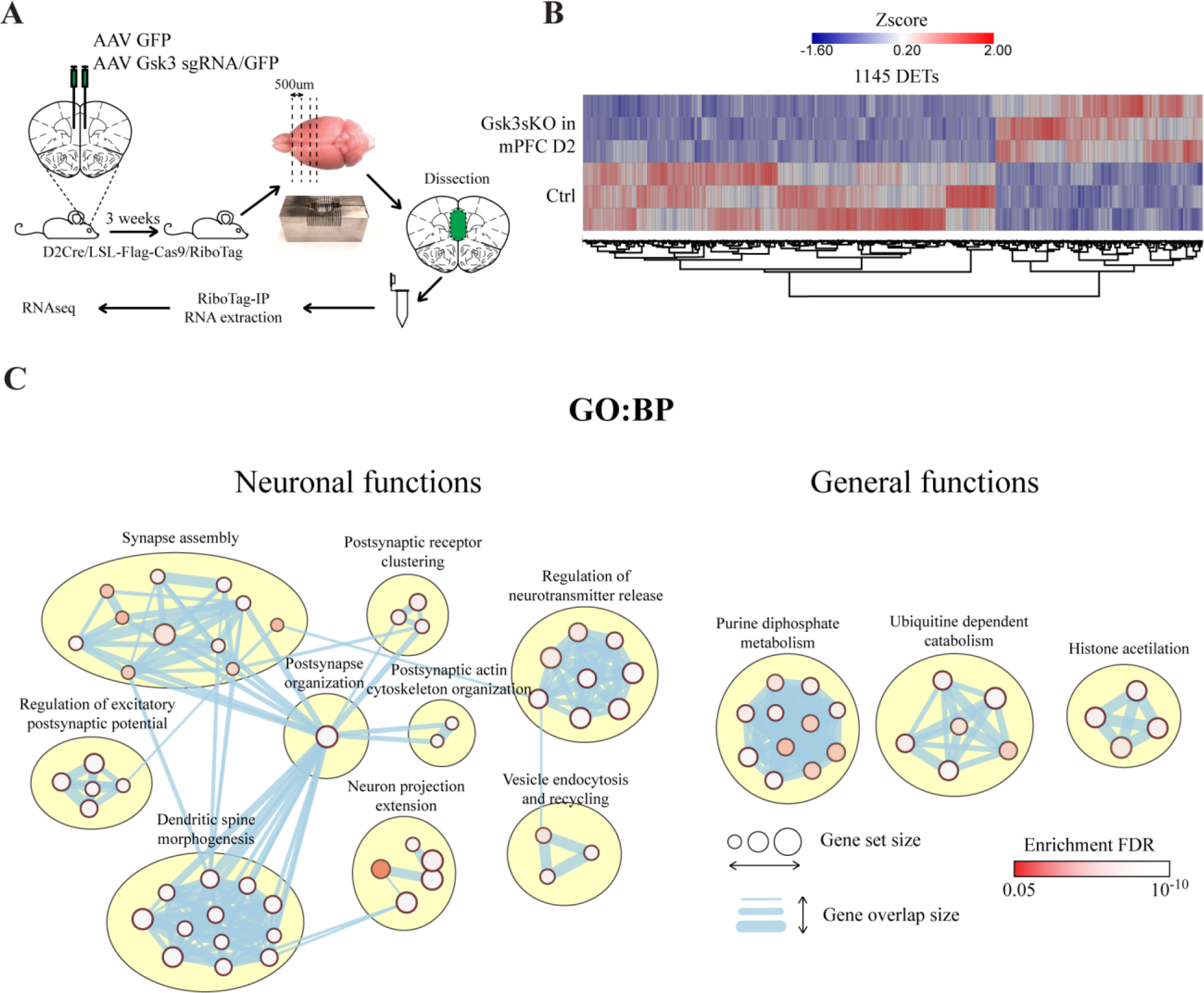
Knockout of GSK3β in D2 neurons of mPFC induces translatome alterations. (A) Schematic representation of virus injection, tissue dissection and RiboTag-IP RNA isolation. (B) Heat map summarizing differentially expressed transcripts (DETs) between Ctrl and Gsk3sKO in mPFC D2 conditions. **c** Enrichment of DETs in GO:BP. Visualization is made by cytoscape.

Overall, this shows that intersectional CRISPR/Cas9 mediated manipulations can be combined with genome wide translational readout. It also indicates that GSK3β knockout in mPFC D2 neurons can induce large scale changes in neuronal translatome.

### Gsk3sKO in mPFC D2 neurons exerts an anxiolytic-like effect on mouse behavior

Behavioral consequences of GSK3β expression in adult D2 expressing mPFC neurons were then investigated. First, Ctrl and Gsk3sKO in mPFC D2 mice were subjected to several anxiety-related behavioral tests. In dark-light emergence test (DLET), Gsk3sKO in mPFC D2 mice spent more time, traveled longer distance as well as performed more entries in the light chamber as compared to control (Figure 4A-D). Open field test (OFT) did not reveal significant differences between the two groups (Figure 4E-G) and in elevated plus maze (EPM) test Gsk3sKO in mPFC D2 mice spent more time on open arms compared to Ctrl mice (Figure 4H). To obtain a more complete understanding of anxiety status we summarized the results across all the tests by performing behavioral Z-scoring for all mice (33, 36). Mice from Gsk3sKO in mPFC D2 group showed a decrease in emotionality Z score compared to control mice (Figure 4I), while their locomotion Z score was unaffected (Figure 4J).

**Figure 4.**
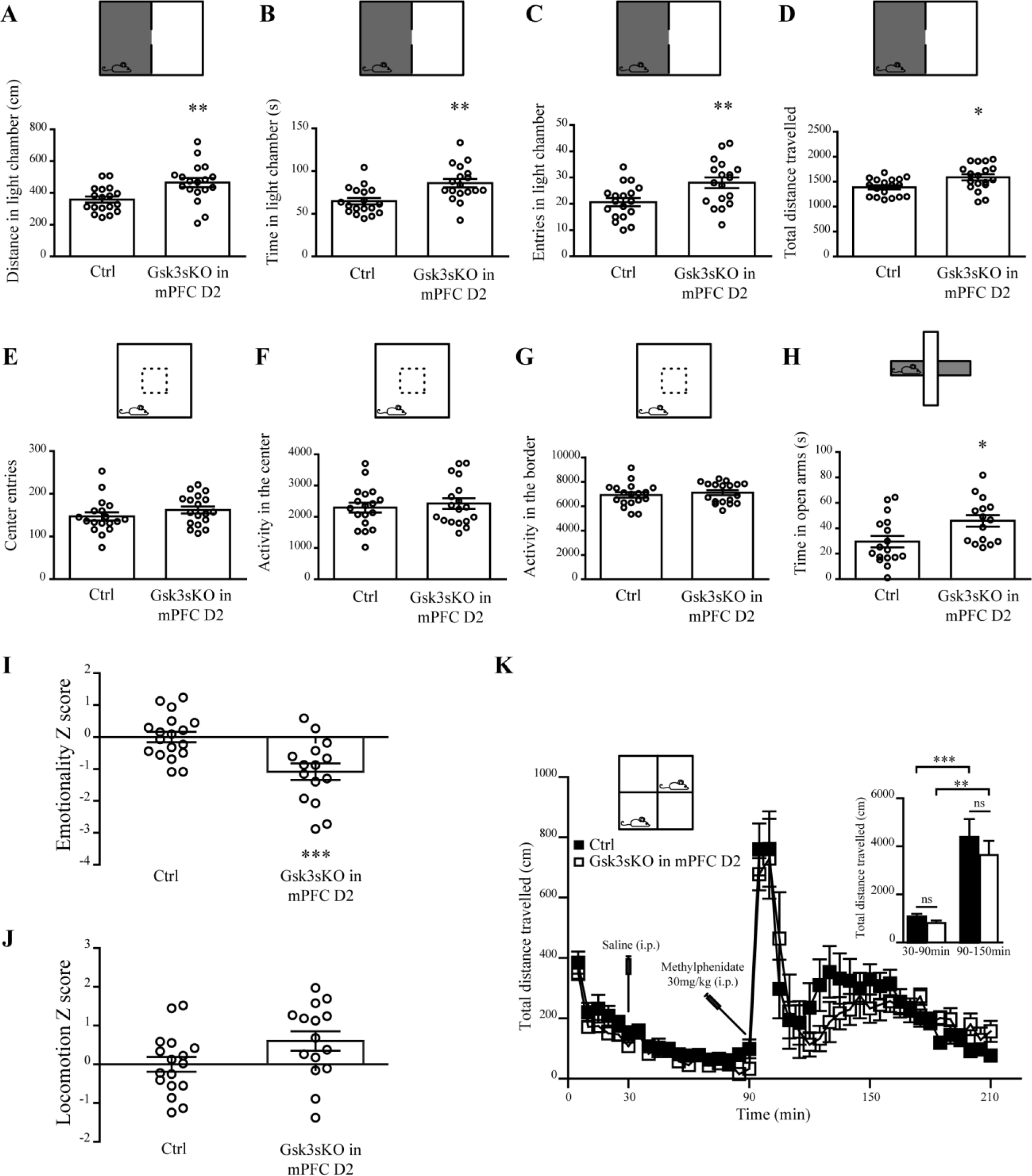
GSK3β of mPFC D2 neurons regulates anxiety-related behaviors. (A-D) Dark/Light emergence test for Ctrl and Gsk3sKO in mPFC D2 mice. (A) Distance travelled in the light chamber (Ctrl: 357.9 ± 18.8cm n=18, Gsk3sKO in mPFC D2: 464.5 ± 29.1cm n=18). (B) Time spent in the light chamber (Ctrl: 64.6 ± 3.7s n=18, Gsk3sKO in mPFC D2: 85.9 ± 4.9s n=18). (C) Entries in the light chamber (Ctrl: 20.6 ± 1.5 n=18, Gsk3sKO in mPFC D2: 28.00 ± 2.0 n=18). (D) Total distance travelled (Ctrl: 1387 ± 41.3cm n=18, Gsk3sKO in mPFC D2: 1585 ± 61.6cm n=18). E-G Open field test for Ctrl and Gsk3sKO in mPFC D2 mice. (E) Center entries (Ctrl: 147 ± 9.5 n=18, Gsk3sKO in mPFC D2: 162 ± 8.3 n=18). (F) Activity in the center (Ctrl: 2293 ± 159.1 n=18, Gsk3sKO in mPFC D2: 2425 ± 170.1 n=18). (G) Activity in the border (Ctrl: 6923 ± 227 n=18, Gsk3sKO in mPFC D2: 7111 ± 198 n=18). (H) Elevated plus maze test for Ctrl and Gsk3sKO in mPFC D2 mice. Time in open arms (Ctrl: 29.5 ± 4.5s n=17, Gsk3sKO in mPFC D2: 45.8 ± 4.5s n=15). (I) Emotionality Z score (Ctrl: −0.0000000033 ± 0.16 n=18, Gsk3sKO in mPFC D2: −1 ± 0.25 n=15). (J) Locomotion Z score (Ctrl: −0.00000001 ± 0.19 n=18, Gsk3sKO in mPFC D2: 0.6 ± 0.25 n=15). (K) Total distance travelled per every 5 min for Ctrl and Gsk3sKO in mPFC D2 mice after injection of Saline at 30 min and Methylphenidate at 90 min. Insert on the right shows total distance travelled between 30-90 min (Ctrl 1067 ± 121.2 n=11, Gsk3sKO in mPFC D2 805.4 ± 105.8 n=10) and 90-150 min (Ctrl 4384 ± 750.0 n=11, Gsk3sKO in mPFC D2 3639 ± 590.8 n=10). Error bars show standard error of the mean (SEM). (*p < 0.05, **p < 0.01, ***p < 0.001, Student T-test).

To evaluate whether GSK3β in mPFC D2 can affect hyperactivity Ctrl and Gsk3sKO in mPFC D2 mice were treated with the dopamine reuptake inhibitor methylphenidate (30mg/kg, i.p.) and locomotion was measured (Figure 4K) before and after drug administration. Mice from both groups showed no difference in exploratory locomotor activity prior to drug administration. Furthermore, in both groups, drug-induced hyperlocomotion was not affected by the knockout of GSK3β in mPFC D2 (Figure 4K).

This indicates that a reduction of GSK3β expression in D2 neurons of mPFC is sufficient to impact anxiety-related behaviors and does not affect locomotor behaviors.

### Gsk3β in mPFC D2 neurons is implicated in cognitive and social behaviors

Cortical GSK3β has been shown to be involved in cognitive and social behaviors (8, 12). Thus, we expanded behavioral testing of both groups of mice to assess working memory and social interactions. In novel object recognition (NOR) test, Gsk3sKO in mPFC D2 mice showed less robust discrimination between novel and familiar objects (Figure 5A-F). This is indicative of mild short-term memory impairments.

**Figure 5.**
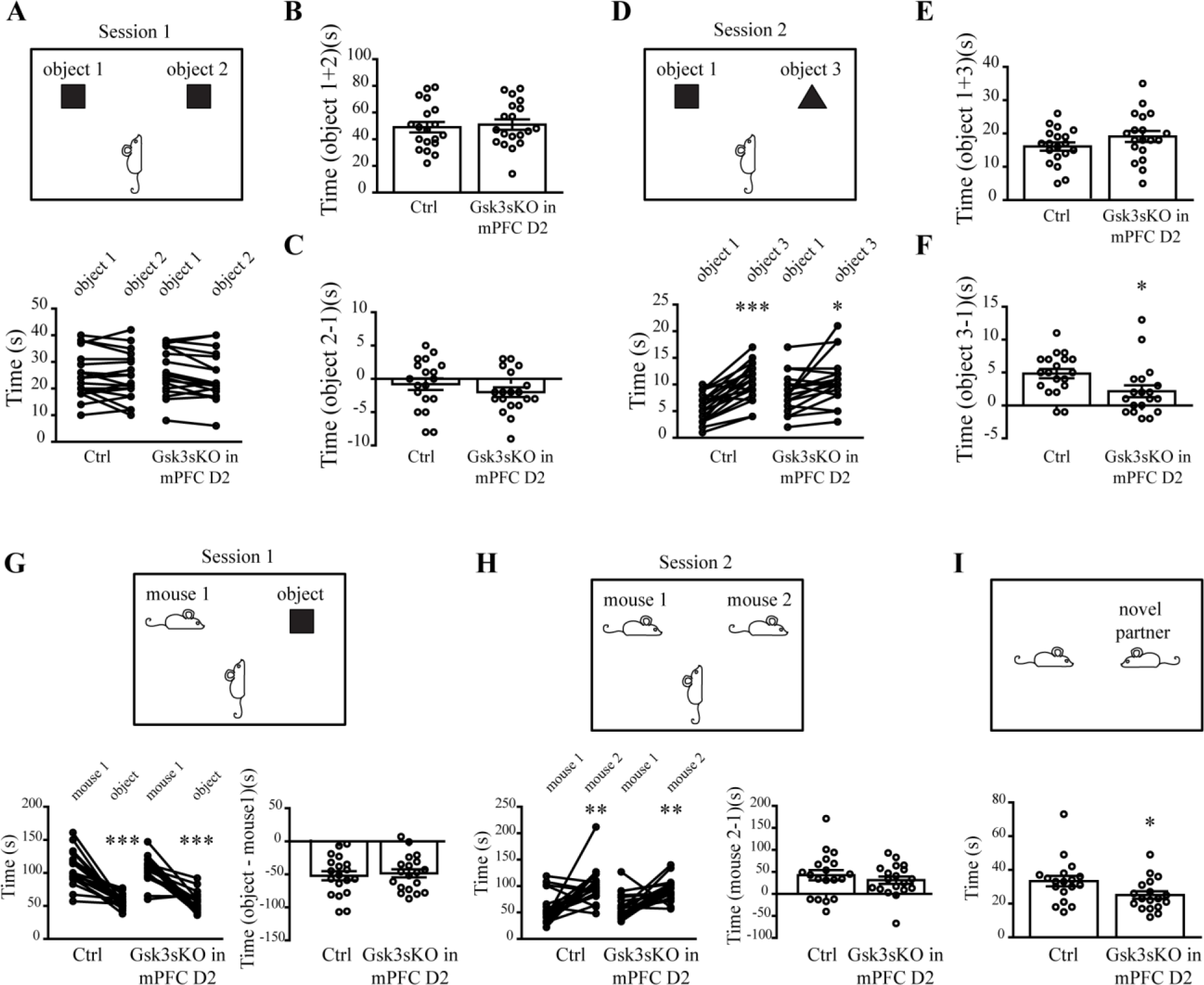
GSK3β of mPFC D2 neurons contributes to cognitive and social behaviors. (A) Novel object recognition test for Ctrl and Gsk3sKO in mPFC D2 mice Session 1. (B) Total time spent with both objects (Session 1 Ctrl: 49 ± 3.9s n=19, Gsk3sKO in mPFC D2: 50.9 ± 3.8s n=19). (C) Difference of time spent with object 2 versus object 1 (Session 1 Ctrl: −0.7 ± 0.8s n=19, Gsk3sKO in mPFC D2: −2 ± 0.7s n=19). (D) Novel object recognition test for Ctrl and Gsk3sKO in mPFC D2 mice Session 2 Session 2 (*p < 0.05, ***p < 0.001, Paired T-test). (E) Total time spent with both objects (Session 2 Ctrl: 16.1 ± 1.2s n=19, Gsk3sKO in mPFC D2: 19.1 ± 1.6s n=19). (F) Difference of time spent with object 3 versus object 1 (Session 2 Ctrl: 4.8 ± 0.7s n=19, Gsk3sKO in mPFC D2: 2.1 ± 0.9s n=19) (*p < 0.05 Student T-test). Preference for social novelty for Ctrl and Gsk3sKO in mPFC D2 mice (G) Session 1 (Time difference object-mouse: Ctrl: −52 ± 6.8s n=19, Gsk3sKO in mPFC D2: −48.4 ± 6.1s n=19), (H) Session 2 (Time difference mouse 2 – mouse 1: Ctrl: 42.5 ± 11.5s n=19, Gsk3sKO in mPFC D2: 31.3 ± 8.3s n=19). (**p < 0.01, ***p < 0.001, Paired T-test). (I) Free social interaction test for Ctrl and Gsk3sKO in mPFC D2 mice. Time in interaction (Ctrl: 33.3 ± 3.1 s n=18, Gsk3sKO in mPFC D2: 25.1 ± 2.2s n=18, *p < 0.05, Student T-test). Error bars show standard error of the mean (SEM).

In the three-chamber social test Gsk3sKO in mPFC D2 mice did not differ from control mice in their preference for social novelty (Figure 5G, H). However, mice in Gsk3sKO in mPFC D2 group spent less time in free social interaction compared to controls (Figure 5I).

Overall, this suggests that GSK3β in mPFC D2 neurons may contribute to short-term memory and social behaviors.

## Discussion

Results presented here underscores the involvement of GSK3β in cortical D2 neurons in different behavioral modalities and demonstrates the usefulness fo CRISPR/Cas9 mediated intersectional knockout strategies to investigate the functions of ubiquitously expressed genes in complex tissues.

D2 receptors are expressed in a mosaic of cell populations across different cortical regions (29). GSK3β is ubiquitously present throughout the brain and is regulating a variety of processes such as neuronal development (2, 3), and synaptic plasticity (5, 6). Among these functions, GSK3β is an important mediator of D2 signaling (24) that can be involved in the effects of lithium and antipsychotic drugs (27, 28). D2 receptor signaling in different brain regions is believed to contribute to different behavioral dimensions under basal conditions and in response to drug treatment (43). Thus, it is required to target GSK3β gene expression only in D2 cells of a given brain region during adulthood if one wants to bypass developmental effects and understand the circuit correlate of its involvement in behavioral regulation. However, this type of genetic manipulation has been technically challenging. We overcame this limitation by combining AAV mediated delivery of *Gsk3b* sgRNA in mice expressing Cas9 only in D2 neurons and achieved efficient and specific high-resolution knockout of GSK3β only in D2 neurons of adult mouse mPFC (Figure 2). One recent study also used an intersectional CRISPR/Cas9 mediated knockout to delete dopamine beta hydroxylase from a defined cluster of tirosine hydroxylase expressing neurons in locus coeruleus to decouple effects of release of norepinephrine from changes of neuronal activity (44). Along with our results this shows that intersectional CRISPR/Cas9 techniques can have a broad applications.

We have investigated whether GSK3β of mPFC D2 neurons is involved in the regulation of anxiety-related, cognitive and social behaviors. These behaviors are known to be affected when manipulating GSK3β activity in a less selective manner (45).

Knockout of GSK3β in mPFC D2 neurons resulted in anxiolytic-like behaviors (Figure 4). In line with our findings, decrease anxiety-related behaviors were documented in forebrain pyramidal neuron GSK3β knockout mice (8), adult PFC GSK3β knockout mice (9) and adult mPFC neurons GSK3β knockout mice (11). However, knockout of GSK3β in all D2 neurons of the brain failed to show anxiety-related effects (28). These differences can be explained by a lack of brain region selectivity and potential developmental effects of deleting GSK3β in all neurons of the brain that expressed D2 either at the adult stage or during development, which can alter the proper formation of neuronal networks. Stimulation of D2 receptor results in an increase of GSK3β activity (45). Interestingly, elevated D2 receptor availability in the prefrontal cortex was found in patients with anxiety disorder compared to healthy subjects (46). Moreover, a decrease in anxiety symptoms after therapy was significantly correlated with the decrease of D2 receptor availability in the prefrontal cortex (47). These findings strongly support a role of GSK3β in the regulation of anxiety-related behaviors downstream of D2 receptors in mPFC of the adult brain.

Knockout of GSK3β in mPFC D2 neurons negatively impacted working memory and free social interaction (Figure 5). Previous reports concerning the involvement of brain GSK3β in the regulation of these behavioral dimensions have been contradictory. In contrast to our findings, gain of function resulting from germline transgenic mice expressing constitutively active GSK3β (Gsk3β KI mice) show impairments in novel object recognition (12). In addition, heterozygous germline GSK3β knockout mice show no difference in social interactions, while forebrain pyramidal neuron GSK3β knockout mice show pro-social effects compared to controls in a free interaction test (8). Differences between these models can once again arise from developmental effects and circuit-specific involvement of GSK3β activity. In line with our observations, antagonism of D2 receptors in the PFC of adult rats impaired novel object recognition and social novelty discrimination (48). This highlights that the involvement of GSK3β in the regulation of cognitive and social behaviors is cell type, brain region and developmental stage selective.

In addition to behavioral changes, the combination of CRISPR/Cas9 knockout with the RiboTag reporter system allowed extracting ribosome associated RNA specifically from mPFC D2 neurons and showed that GSK3β KO has large impact on neuronal translatome (Figure 3). Particularly, transcripts affected by GSK3β KO were enriched in synaptic structure and function, neurotransmission, purine metabolism, ubiquitine dependent catabolism and histone acetylation among others. This underscores the possibility of obtaining translatome footprint of a given gene knockout in a brain region and cell type selective manner.

Inhibition of GSK3β activity has been reported to occur in response to SSRIs, ketamine, antipsychotics, some anticonvulsants (mood stabilizers) and lithium (9, 14, 15, 25−27). Furthermore, data from animal models support a role for GSK3β activity in the effect of these drugs (45). It is probable that subsets of these effects are explainable by modulation of GSK3β downstream of D2 receptors in different brain regions. Our results indicate that GSK3β in adult mPFC neurons expressing D2 contributes to cognitive, social and emotional dimensions of behavioral regulation. This regulation can for example contribute to the regulation of mood by lithium or of negative symptoms by antipsychotics in schizophrenia.

Intersectional approaches have proven important to the precision of functional studies of neuronal circuits using optogenetics. Our results underscore the viability of this approach to also study gene functions in a network defined fashion. This can advance our understanding of drug action at a brain circuit level and potentially lead to the development of circuit selective therapeutics.

## Funding and Disclosure

This work was supported by an operating grant from the Canadian Institutes of Medical Research (CIHR) MOP-136916.

## Author contribution

JK and J-MB conceived the study, designed the experiments and wrote the paper. JK performed the experiments and analyzed the data.

## Additional information

### Competing interests

The authors declare no competing interests.

