## Supplementary Material for "CRISPR/Cas9 mediated intersectional knockout of GSK3β in D2 receptor expressing mPFC neurons reveals contributions to emotional regulation"

### Supplementary Materials and Methods

#### Cell line culture, transfection, western blot and TIDE analysis

Neuro-2A cells were grown in high glucose DMEM containing 10% FBS, penicillin/streptomycin and L-glutamine (HyClone-GE Healthcare, Logan, UT). Cells were maintained at 37°C in 5% CO2 atmosphere and transfected using Lipofectamine 2000 (Thermo Fisher Scientific, Waltham, MA) according to the manufacturer's protocols.

For TIDE analysis and in vitro evaluation of Gsk3β and Gsk3α expression by Western blot (Fig. 1), 50–70% confluent N2A cells were transfected with previously validated all in one px459 based constructs (pX459 vectors with guide targeting *Gsk3b*) (1). To select only transfected cells, 48 hours after transfection cells were incubated with 3µM puromycin for 72 hours followed by 48 hours incubation without puromycin.

For western blot, cells were washed and lysed on the day 7 after in lysis buffer containing: 50mM Tris-HCl, 150mM NaCl, 5mM EDTA, Protease inhibitor cocktail, 1% SDS, 0.5% Na-deoxycholate, 1% NP-40, 10mM NaFluoride, 25mM βglycerophosphate, 10mM Na Orthovanadate (Sigma-Aldrich, Oakville, Canada).  Lysates were centrifuged 10000g for 30 min and supernatants were collected. Protein concentration was measured by using a DC-protein assay (Bio-Rad, Hercules, CA). Protein extracts were separated on precast 4-20% Tris-glycine gels (Thermo Fisher Scientific, Waltham, MA) and transferred to nitrocellulose membranes. Blots were immunostained overnight at 4 °C with primary antibodies. Immune complexes were revealed using appropriate IR dye-labeled secondary antibodies from Li-Cor Biotechnology (Lincoln, NE). Quantitative analyses of fluorescent IR dye signal were carried out using Odyssey Imager and Image Studio Lite 5.2 software (Licor Biotechnology, Lincoln, NE). For quantification, GAPDH was used as a loading control for the evaluation of total protein levels.

Following primary antibodies were used in the experiments: mouse anti-GAPDH (1:5000, Santa Cruz sc-322333 ), mouse anti-Gsk3α/β (1:500, Santa Cruz sc-7291). Secondary antibodies: goat anti-mouse IR Dye 680 (1:10000, Mandel 926-68020).

To isolate genomic DNA for TIDE analysis (2), cells were lysed by tail buffer (Tris pH=8.0 0.1M, NaCl 0.2M, EDTA 5mM, SDS 0.4% and proteinase K 0.2mg/ml), and DNA was precipitated using isopropanol followed by centrifugation (13000g 15min). DNA was resuspended in TE Buffer (10 mM Tris pH 8.0, 0.1 mM EDTA) and used for PCR. Following primers were used to amplify On-Target and 3 putative Off-Target sites:

| Genomic region | Forward primer sequence | Reverse primer sequence |
|---|---|---|
| On-Target site | GGTTCCTCTTGCCCCCTATTA | TTCTCATTGGCATTTCCACGC |
| Off-Target site 1 | TCATTATAGGTCTCGGGGCAAG | AAATGATGAAGGAATTTGGTCGGAA |
| Off-Target site 2 | CCTGCTGTCTCTCCCTTGTG | GGAAGCCTACTGCAAGAGCA |
| Off-Target site 3 | AACGTGAAACTTTGTTTGCAATATC | ACAAATTTCAATCTGTGGCTGGG |

PCR products were sent to sequencing with Forward primers and frequencies of mutations were determined by online TIDE tool (https://tide.nki.nl/) (2).

### Behavioral tests

#### Open field test (OFT)

OFT was performed for 30 min in an automated Omnitech Digiscan apparatus (AccuScan Instrument, Columbus, OH). Each mouse was placed in a corner of the large Plexiglas box and the exploratory activity was recorded. A number of entries, time, horizontal activity and total distance were recorded separately for the central (25% of the total surface) and peripheral areas.

**Dark-light emergence test (DLET)**

DLET was performed for 5 min with mice placed initially at the center of the dark chamber. Tests were conducted using an automated open field activity apparatus with light/dark insert (Med-Associates, St Albans, VE) with the light compartment illuminated at 800 lux. The total time spent in the dark and light compartments, the total distance traveled, and the number of entries from the dark to the light chamber was used as parameters for analysis.

**Elevated plus maze (EPM)**

EPM was performed for 5 min with mice initially placed in the far end of the close arm. Mice were video tracked using Viewer software (Biobserve behavioral research). The time spent in the open arm was measured and used for the analysis.

**Behavioral Z scoring**

To obtain integrated measures in each group, emotionality- and locomotion-related data were normalized using a Z-score methodology (3). Z-scores for individual animals were calculated using the formula: $Z = (X-\mu)/\sigma$, which indicates how many standard deviations ($\sigma$) an observation ($X$) is above or below the mean of a control group ($\mu$). Z-scores for behavioral measures were first averaged within the test, and then across all three tests (OFT, DLET, EPM). OFT (center entries), DLET (distance in the light chamber), EPM (time in open arms) were used to obtain emotionality Z-scores. Locomotion Z-scores were obtained from DLET (total distance traveled) and OFT (distance traveled in the border) data.

**Novel object recognition (NOR)**

The test was performed in the box with dimensions measuring 60x40x20 (length, width, height). Mice were habituated to the arena for 5 min 24hours before the test. The test consisted of 2 sessions. On session 1, mice were allowed to explore 2 identical objects placed in the left and the right side of the arena for 10 minutes. For session 2, 1 hour after session 1, one of the familiar objects was changed with a novel object and mice were allowed to explore for 5 min. Objects we thoroughly cleaned between the sessions and between different mice. All the process was video recorded and the time that mice investigated (sniffed) the objects were measured manually (by the observer being unaware of the treatment).

**Preference for social novelty**

The test was performed in the box with dimensions measuring 60x40x20 (length, width, height) that was separated into 3 equal compartments with a clear walled Plexiglas dividers with doors, allowing test mice to freely explore all 3 compartments. Each outer compartment also contained an inverted cup (Galaxy Pencil cup/Utility Cup; Spectrum Diversified Designs, Inc., Streetsboro, OH, USA). Before the beginning of the test, mice were placed in the box for 5 min to habituate. The test consisted of 2 sessions. For session 1, an unfamiliar mouse was placed in the left inverted cup and an object was placed in the right inverted cup. Then the test mouse was free to explore all 3 compartments for 10 minutes. For session 2, the object in the right cup was replaced by a novel mouse. Again the test mouse was free to explore all 3 compartments for 10 minutes. All the process was video recorded and the time that the test mouse investigated object or mice in the cups were measured manually (by the observer being unaware of the treatment).

**Free social interaction**

The social interaction test was performed in a clear Plexiglas box, with dimensions measuring 30x30x20 cm (length, width, height). Mice were first habituated to the arena for 5 min 24 hours before the test. On the day of the test, subject mice were paired with age and sex-matched unfamiliar mice from the D2Cre/LSL-Flag-Cas9 colony for 10 minutes. The entire process was video recorded and the time spent in social interaction (sniffing and grooming the partner) was measured manually (by the observer being unaware of the treatment).

**Table S1**. **Differentially expressed transcripts (DETs) between Ctrl and Gsk3sKO in D2 conditions**

Shows differentially expressed transcripts in Gsk3sKO in D2 compared to Ctrl mice. DETs with $p<0.05$ are shown.

**Table S2. gProfiler enrichment of DETs in biological pathways**

Shows significantly enriched pathways, enrichment FDRs and the list of enriched transcripts.

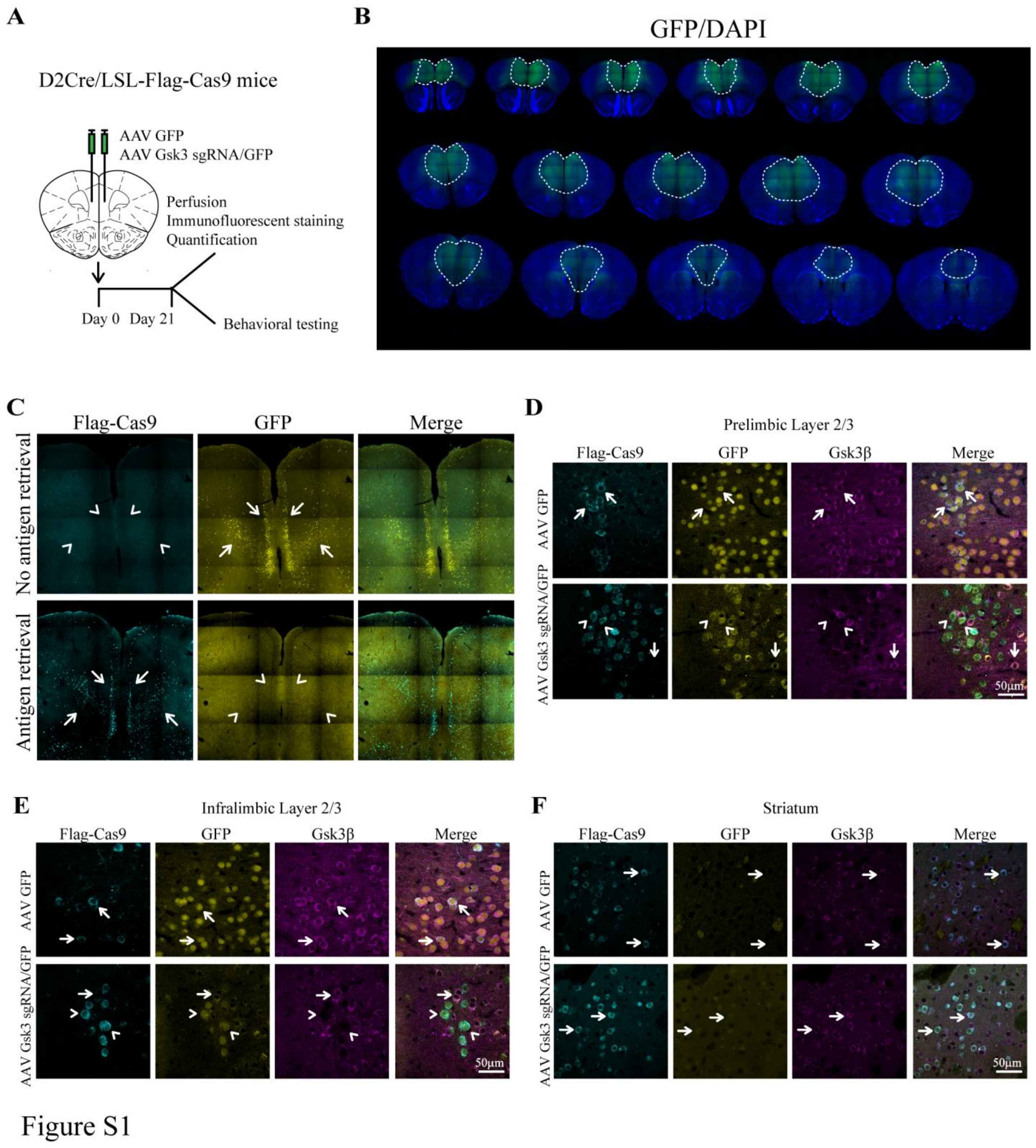


**Figure S1. Cas9 expression and knockout of Gsk3β in mPFC D2 neurons.**

(A) Schematic representation of stereotaxic injection of viruses and experimental design.

(B) Serial coronal sections showing the infection volume of the AAV GFP and AAV Gsk3 sgRNA/GFP viruses. (C) Flag-Cas9 staining is detected after antigen retrieval in mPFC. Endogenous GFP signal in D2Cre/LSL-Flag-Cas9 mice disappears after antigen retrieval. Arrows show presence and arrowheads show absence of fluorescent signal. (D-F) Immunofluorescent staining for Flag-Cas9 and Gsk3β in (D) Prelimbic layer 2,3, (E) Infralimbic layer 2,3, (F) Striatum of virus injected D2Cre/LSL-Flag-Cas9 mice. Note that in AAV GFP injected control condition all cells express Gsk3β (indicated by arrows). In AAV Gsk3 sgRNA/GFP injected condition only cells that express Flag-Cas9 (corresponding to D2 cells) and sgRNA/GFP do not have Gsk3β signal (indicated by arrowheads), while cells having sgRNA/GFP, but not Flag-Cas9 staining (not D2 cells) still express Gsk3β (indicated with arrows). In Striatum all the cells in both conditions express Gsk3β, since the viruses were injected into mPFC and did not infect Striatum.

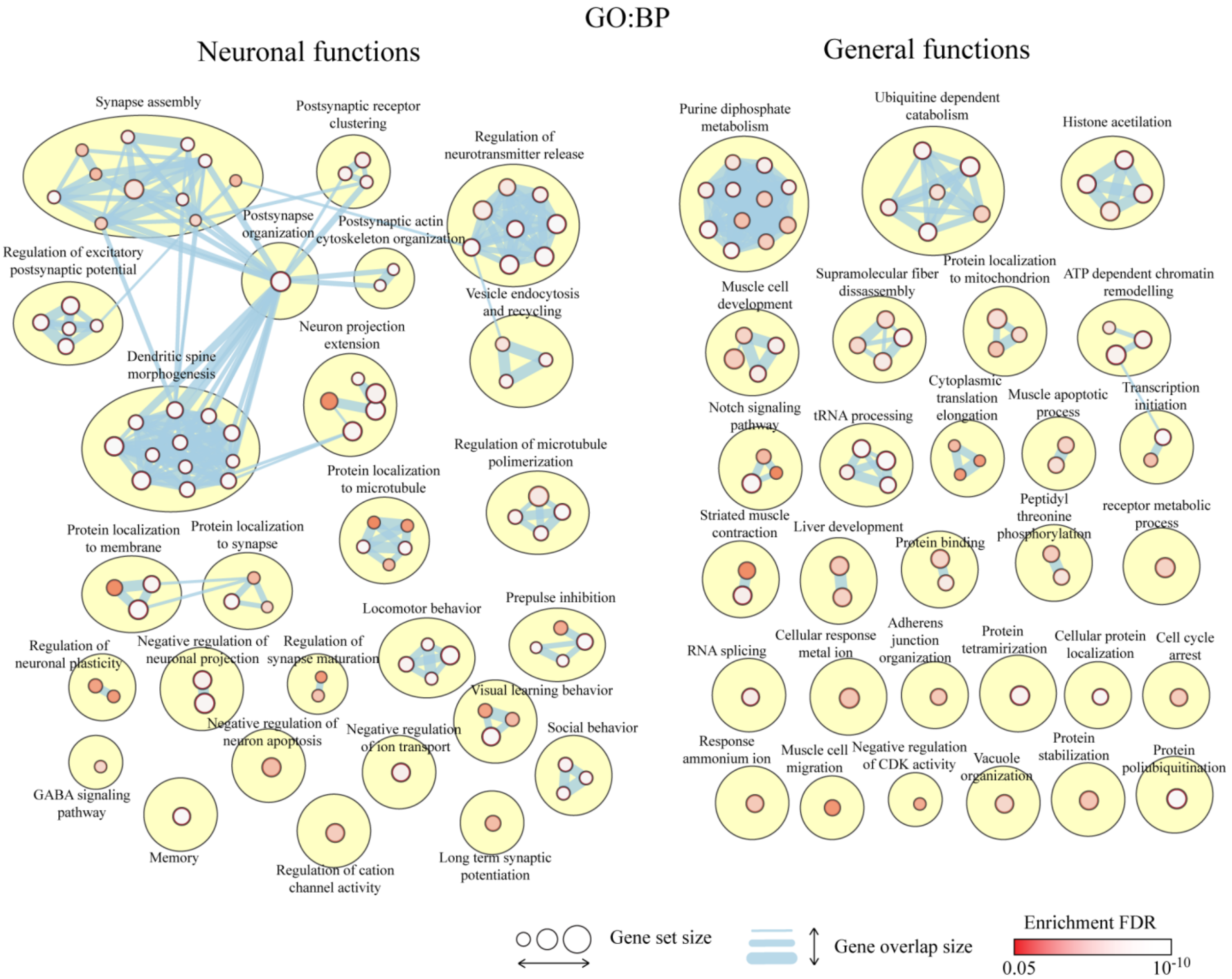


Figure S2

**Figure S2. Enrichment of DETs in biological pathways**

Enrichment of differentially expressed transcripts (DETs) between Ctrl and Gsk3sKO in mPFC D2 in GO:BP. Visualization is made by cytoscape.

1. Khlghatyan J, Evstratova A, Chamberland S, Marakhovskaia A, Bahremand A, Toth K, et al. (2018): Mental Illnesses-Associated Fxr1 and Its Negative Regulator Gsk3β Are Modulators of Anxiety and Glutamatergic Neurotransmission. *Front Mol Neurosci*. 11:119.
2. Brinkman EK, Chen T, Amendola M, van Steensel B (2014): Easy quantitative assessment of genome editing by sequence trace decomposition. *Nucleic Acids Res*. 42:e168.
3. Guilloux JP, Seney M, Edgar N, Sibille E (2011): Integrated behavioral z-scoring increases the sensitivity and reliability of behavioral phenotyping in mice: relevance to emotionality and sex. *J Neurosci Methods*. 197:21-31.
